# Limited genetic diversity among *Malassezia furfur* isolates during two one-year neonatal and pediatric bloodstream infection surveys confirms preference for one specific IGS1 genotype in deep-seated infections

**DOI:** 10.1101/2025.01.20.633816

**Authors:** Bart Theelen, Claudia Cafarchia, Ferry Hagen, Teun Boekhout

**Author notes:** During part of this work at Westerdijk Fungal Biodiversity Institute, Utrecht, Netherlands (Teun Boekhout).

## Abstract

In recent years, multiple reports indicated that *Malassezia* fungemia may be of emerging concern in immunocompromised patients and neonates. In combination with observations of reduced antifungal susceptibility, this highlights the need for a better understanding of the extent of *Malassezia* etiology in deep seated infections. The use of media without lipid supplementation in most clinics underlines the likelihood of underdiagnosis. Here, we utilize multiple genetic markers to explore the genetic diversity of number one *Malassezia* fungemia species *Malassezia furfur*, observed from two one-year surveys investigating neonatal and pediatric bloodstream infections. All strains isolated from skin and deep seated neonatal or pediatric sites belong to IGS1-genotype A, except for nine isolates from one pediatric patient. This confirms previous observations of a preferential genotype for deep seated body sites. The mitochondrial *cox*3-*nad*3 intergenic region provided additional typing resolution, and including ITS, translation elongation factor 1-α, chitin synthase 2, and the mating type, 16 genotypes (MF1-16) could be distinguished among all isolates from both surveys and reference isolates. These results present a promising MLST scheme that requires further testing. We observed a decreased genetic variation of *M. furfur* isolates among NICU and pediatric surgery patients over the course of 5 years and showed that the skin likely serves as a reservoir for resident *M. furfur* strains from where they can enter the bloodstream, causing infections among susceptible patient groups. Overall, reduced susceptibility for amphotericin B and specifically fluconazole was observed.

## Introduction

*Malassezia*, the most abundant fungal genus on healthy human skin (Findley et al., 2013), takes on a pathogenic role in various skin disorders, including seborrheic dermatitis (SD) / dandruff (D), pityriasis versicolor (PV), atopic dermatitis (AD) and *Malassezia* folliculitis (Ianiri et al., 2022; Saunte et al., 2020). In recent years, *Malassezia* has attracted increased attention for its involvement in diseases such as Crohn’s disease (Limon et al., 2019) and pancreatic cancer (Aykut et al., 2019), illustrating colonization of deeper body sites or transmission from skin to these sites with unfavorable health effects.

A more obvious transmission is that from skin to bloodstream in situations when the skin barrier is disrupted, for example when intravenous lipid-rich nutrition is administered in immunocompromised patients or neonates (Rhimi, et al., 2020). Multiple cases have been described over the years but the extent of *Malassezia* involvement in bloodstream infections (BSIs) is unknown (Priputnevich 2018; Rhimi et al. 2020; Chow et al. 2020; Huang et al. 2020). Due to the lipid dependence of *Malassezia* yeasts and the use of standard culture media without lipid supplementation in most clinics, it has been argued that *Malassezia* BSIs are an underdiagnosed phenomenon (Rhimi, et al., 2020). Only three of the 18 currently described *Malassezia* species have been implicated in BSIs(Rhimi, et al., 2020). Most frequently involved species is *Malassezia furfur*, followed by *Malassezia pachydermatis* which generally is not a colonizer of humans but rather of pet animals such as dogs, and it has been suggested that transfer from pet to owner and healthcare worker are transmission routes to the bloodstream in susceptible patients (Chang et al., 1998; Guillot & Bond, 2020). Only a few cases of *Malassezia sympodialis* BSI have been described to date (Rhimi, et al., 2020).

Most *Malassezia* BSI reports have identified its involvement only to the species level and little is known about the genetic intraspecies variation. One study investigated the molecular epidemiology of a *M. pachydermatis* outbreak in a neonatal unit with IGS1 genotyping, identifying the involvement of multiple genotypes, sometimes even within one patient (Ilahi et al., 2018). Another study, also focusing on *M. pachydermatis*, applied a whole genome-based SNP-analysis, showcasing its potential for outbreak detection (Chow et al., 2020). This species is generally not known to colonize humans. However, this is the only known *Malassezia* species that is able to grow on standard Sabouraud dextrose agar (SDA), which increases the odds for it to be detected in routine clinical procedures (Wu et al., 2015).

*Malassezia furfur*, the number one *Malassezia* BSI species, has been isolated from both humans and multiple animal species, and does require lipid supplementation for successful growth on culture media (Theelen et al., 2018). One study of catheter-associated *M. furfur* from hospitalized adults utilized ITS1-sequencing and showed that all catheter-associated isolates belonged to the same ITS1-genotype. Interestingly, for two patients some of the *M. furfur* isolates taken from the armpit also belonged to that same genotype, suggesting possible transmission from skin to catheter and blood (Kaneko et al., 2012). Some general studies included a few *M. furfur* isolates from deep-seated body sites and it was suggested that deep-seated *M. furfur* isolates preferentially may belong to one unique genotype (Theelen et al., 2001, 2021). Recently it was proposed that certain mitochondrial loci might present excellent targets for typing in *M. furfur* and the *cox*3–*nad*3 mt intergenic region was identified as the most promising mitochondrial target for typing, but only a limited number of isolates from deep-seated sources were included in the evaluation (Theelen et al., 2021).

Two one-year yeast fungemia surveys held in Italy in 2011 and 2016 explored the prevalence of *Malassezia* in BSIs of neonates and pediatric patients and *M. furfur* was identified as the causative species (Iatta et al., 2014; Iatta et al., 2018). One of them also discussed detection and diagnosis considerations of *M. furfur* bloodstream infections (Iatta et al., 2018) and a few studies addressed antifungal susceptibility (AFS) trends also comparing various testing methods (Iatta et al. 2014; Iatta et al. 2015; Rhimi et al. 2020). The above-mentioned studies considered the discussed *M. furfur* isolates as groups and did not look at potential differences between isolates from individual patients or sampling year, and identification was assessed at the species level. Here we explored 125 isolates (including four *M. slooffiae* isolates) obtained from above-mentioned surveys in more detail, together with 13 additional reference isolates derived from various deep-seated sources and geographical locations. We evaluated the validity of previous claims regarding limited genetic variation of deep-seated *M. furfur* isolates, looked at the applicability of various loci for typing, and explored variation between the different surveys, patient groups and sources, in comparison to the reference deep-seated isolates and isolates derived from other sources and belonging to other genotypes. Finally, we evaluated the potential of selected loci for an MLST-scheme in *M. furfur*, as is widely used in pathogenic species such as *Cryptococcus neoformans* and *Candia albicans* (https://mlst.mycologylab.org/, https://pubmlst.org/organisms/candida-albicans). Available antifungal susceptibility (AFS)-data of a subset of isolates were evaluated in correlation with genotypic and source information.

## Results and Discussion

### Italian neonatal and pediatric *M. furfur* isolates confirm a preferential IGS1-genotype for deep seated infections

Previous studies have utilized the intergenic spacer 1 region (IGS1) as a suitable marker for genotyping in *Malassezia restricta* (Sugita et al., 2004) and *Malassezia globosa* (Sugita et al., 2003), and another investigation recently identified six main IGS1 clusters in *M. furfur* with some additional variation among the main groups (Theelen et al., 2021). Previous observations applying Amplified Fragment Length Polymorphism (AFLP) suggested the preference for two related genotypes from deep seated sources (Theelen et al., 2001), which was corroborated for IGS1, but based on only 10 isolates. Deep seated isolates almost exclusively belonged to genotype IGS1-A, with minor subdivision between IGS1-A1 and IGS1-A2 subgenotypes. Here, we further tested these initial findings with 125 clinical isolates from two 1-year *Malassezia* fungemia surveys in neonatal and pediatric wards from Bari, Italy, and 13 additional deep seated reference isolates from various backgrounds and geographies. Of the 139 tested isolates, all belonged to either genotype IGS1-A1 or IGS1A-2, except for nine isolates that all were derived from one pediatric patient and retained genotype IGS1-G (Fig. 1, Table S1). A Russian study between 2015 and 2017, evaluated deep seated *M. furfur* infections for 2793 NICU and surgery NICU patients. In 14 % of surgery NICU patients, *M. furfur* was detected (Priputnevich, 2018). A randomized subset of 94 *M. furfur* isolates was further studied and all belonged to genotype IGS1-A1 (personal observation by B. Theelen). Genotypes IGS1-A1 and IGS1-A2 are very similar, only differing by one SNP. Besides the current work, only one other study was published evaluating sub-species level genetic variation between *M. furfur* isolates from blood, catheter tips, and surrounding skin of hospitalized patients in Japan, but based on the ITS1 region of the ribosomal DNA. The authors identified that all isolates from blood and catheter tips were the same ITS1-type (I-3) (Kaneko et al., 2012). Two of these isolates, GIFU01 and GIFU02, were also included in the current study as reference isolates, and our analysis identified them as IGS1-A2. Taken together, these results confirmed the propensity of a unique IGS1 genotype of *M. furfur* for deep seated infections, in neonates as well as in hospitalized adults, across a wide geographical area and time. The fact that from one patient (patient 4) a different IGS1 genotype was observed, illustrated that this propensity is not absolute but may, to some extent, be context-dependent, in line with the more general observed complexity of *Malassezia* interactions with its environment and host (Vijaya Chandra et al., 2020; Ianiri et al., 2022).

**Figure 1.**
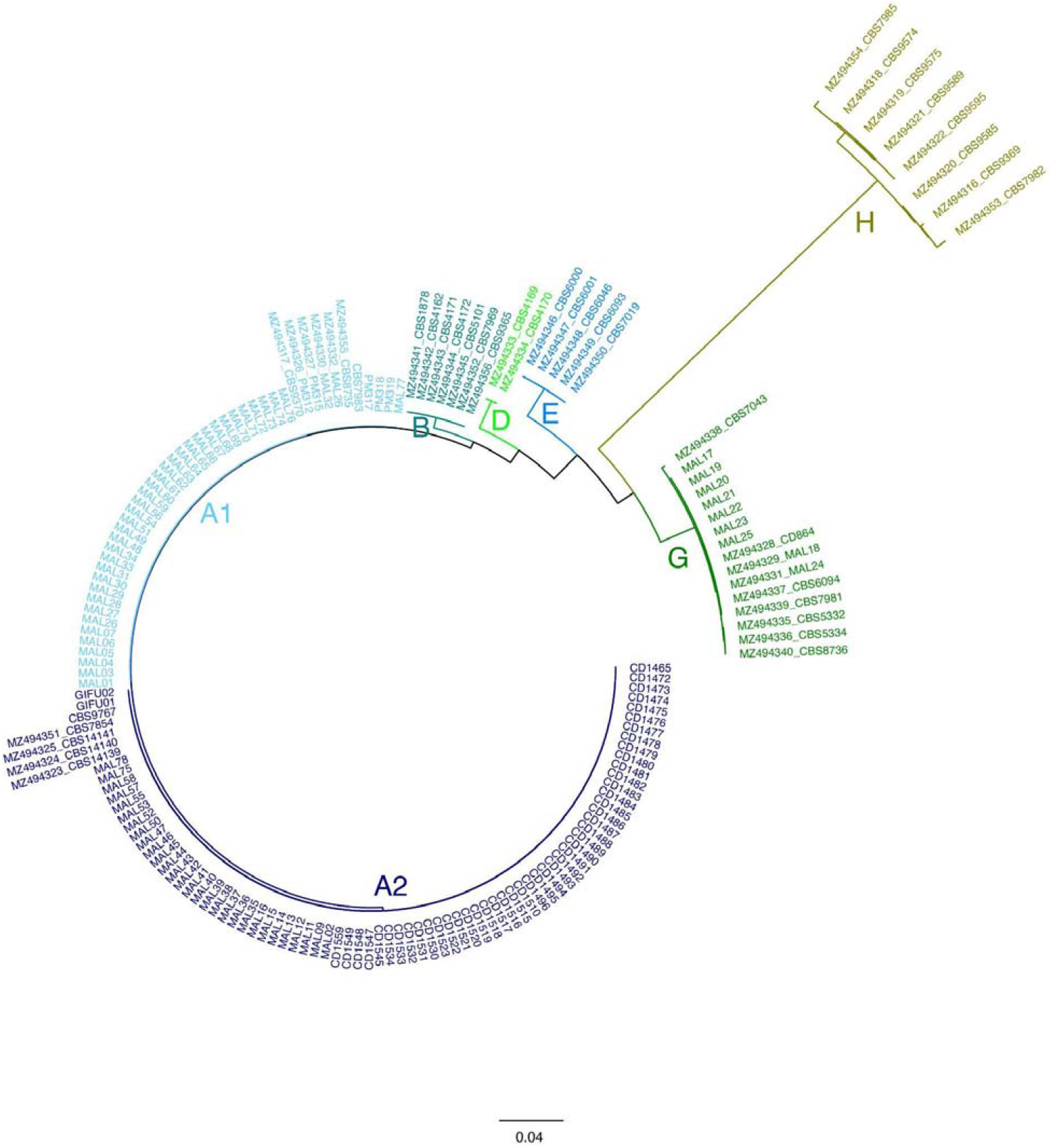
Phylogenetic tree for *Malassezia furfur* based on the intergenic spacer 1 of the ribosomal DNA, for 121 isolates obtained from two one-year surveys in an Italian hospital and 13 reference strains. Clades are labeled with IGS1-genotype codes. Each color represents a genotype.

### Applicability of the mitochondrial *cox*3-*nad*3 intergenic region for *M. furfur* typing

A recent comparative analysis of 20 *M. furfur* mitochondrial genomes identified three variable regions as promising targets for strain typing. A total of 41 strains, representing the six main IGS1-clusters, were sequenced for these three regions and the *cox*3-*nad*3 region was identified as the most promising of the three (Theelen et al., 2021). We evaluated the sequence variation of this region for the 125 Italian isolates and reference isolates and compared these with the previously published 41 strains with few strains overlapping between both studies. Although most isolates from the current study were all identical (i.e., genotype i4), additional genotypes were observed, totaling nine genotypes among the isolates of this study (Fig. 2, Table 1, Table S1). Noteworthy, the nine isolates from one pediatric patient with deviating genotype IGS1-G, also retained a unique genotype for *cox*3-*nad*3 (l7). While the two Japanese isolates from blood and catheter did not differ from the other isolates in this study for IGS1, they carried a unique *cox*3-*nad*3 genotype i3 that was not observed among the 125 Italian isolates or any of the other reference isolates. It is tempting to speculate whether genotype i3 could represent a geographically linked genotype, but this needs to be evaluated with a larger set of isolates. Though this study focuses on isolates from Italian neonatal and pediatric patients, with the inclusion of the previously published *cox*3-*nad*3 sequences (Theelen et al., 2021), the added value of this region for typing is further confirmed.

**Figure 2.**
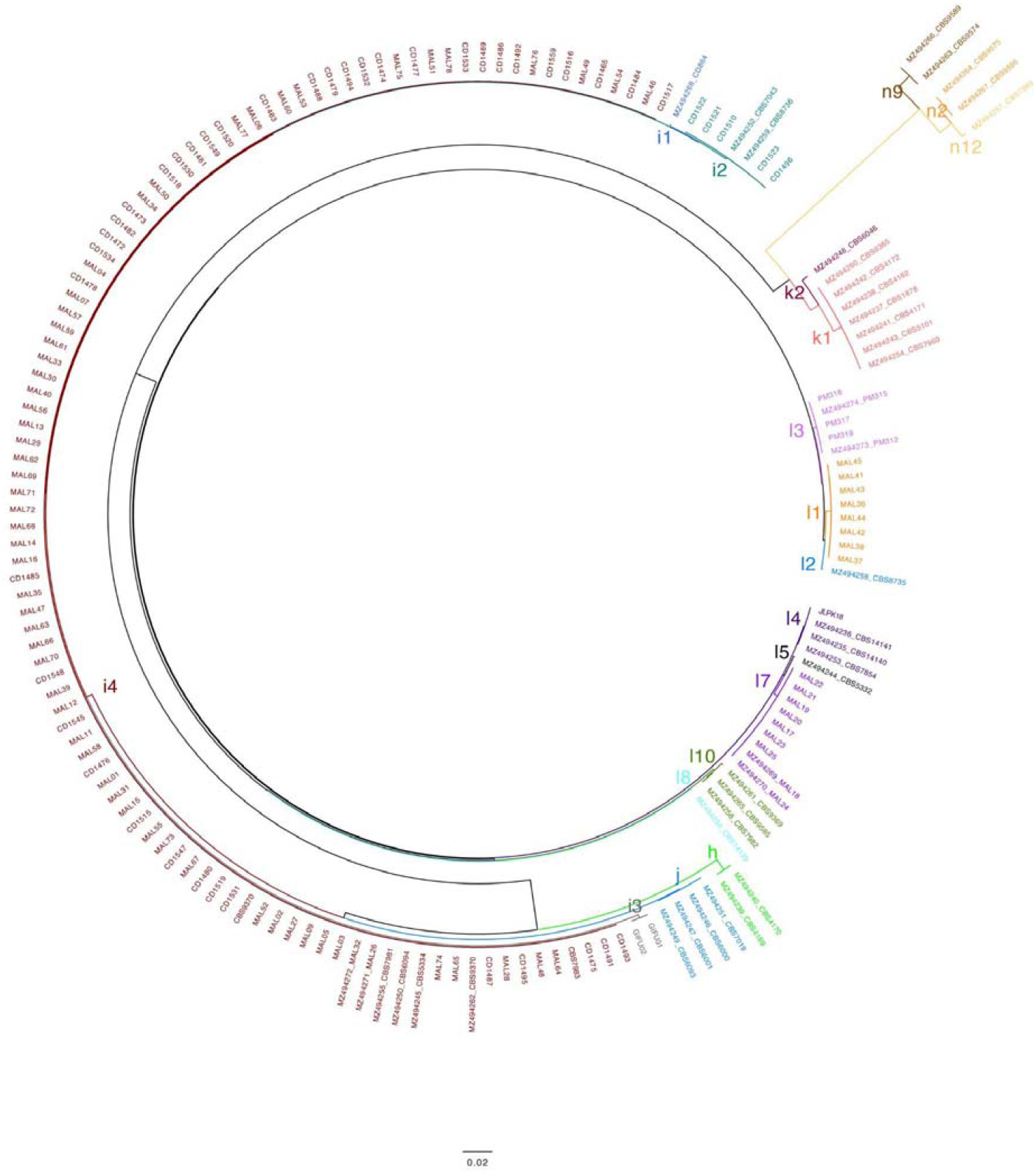
Phylogenetic tree for *Malassezia furfur* based on the *cox*3-*nad*3 intergenic region, for 121 isolates obtained from two one-year surveys in an Italian hospital and 13 reference strains. Clades are labeled with *cox*3-*nad*3-genotype codes. Each color represents a genotype.

**Table 1.** *Malassezia furfur* overall genotypes of strains from two one-year surveys in an Italian hospital and references, based on a combination of five nuclear loci, one mitochondrial locus, and mating type. Abbreviations: MF, *Malassezia furfur*; IGS1, intergenic spacer 1 region; ITS, internal transcribed spacer; Btub, □-tubulin; CHS2, chitin synthase 2; *TEF1*-α, translation elongation factor 1-α, *cox*3-*nad*3, *cox*3-*nad*3 intergenic region.

| MF genotype | IGS1 | ITS | *TEF1*-α | *CHS2* | Btub | *cox3-nad3* | mating type |
| --- | --- | --- | --- | --- | --- | --- | --- |
| MF1 | A1 | K | V | II | $\mu$ | i4 | a1b1 |
| MF2 | A1 | K | V | II | $\mu$ | l2 | a1b1 |
| MF3 | A1 | K | V | II | $\mu$ | l3 | a1b1 |
| MF4 | A1 | K | W | II | $\mu$ | i4 | a2b1 |
| MF5 | A1 | N | V | II | $\mu$ | i4 | a2b1 |
| MF6 | A1 | N | W | II | $\mu$ | i4 | a2b1 |
| MF7 | A2 | N | V | II | $\mu$ | i4 | a1a2b1 |
| MF8 | A2 | N | V | II | $\mu$ | i4 | a2b1 |
| MF9 | A2 | N | V | II | $\mu$ | l4 | a2b1 |
| MF10 | A2 | N | V | II | $\mu$ | l8 | a1b1 |
| MF11 | A2 | N | W | II | $\mu$ | i2 | a2b1 |
| MF12 | A2 | N | W | II | $\mu$ | i4 | a1b1 |
| MF13 | A2 | N | W | II | $\mu$ | i4 | a2b1 |
| MF14 | A2 | N | W | II | $\mu$ | l1 | a1b1 |
| MF15 | A2 | N | W | III | $\mu$ | i3 | a1b1 |
| MF16 | G | L | W | III | $\Delta 2$ | l7 | a2b1* |

**a2b1\*** The “b1” locus has a deviating sequence.

### An MLST scheme test case for *M. furfur* identifies 15 unique genotypes among deep seated isolates

In addition to IGS1 and *COX*3-*NAD*3, four additional loci were included to assess the genotypic variation among the studied isolates, viz., ITS and three protein coding genes □-tubulin (Btub), chitin synthase (*CHS*2), and translation elongation factor 1-α (*TEF1*-α) were included as these are common barcodes used in species description within the genus *Malassezia* and beyond, and they were also found useful for delineating various hybrid and parental clusters within *M. furfur* (Theelen et al., 2022). The mating type was included as an additional marker. Among the 139 tested isolates, when combining all markers, 16 overall *Malassezia furfur* (MF) genotypes were identified, of which only MF11 was not associated with a deep-seated source but found from the ear of a patient (Table 1, Table S1). Except for the mitochondrial barcode *cox*3-*nad*3 (nine observed genotypes), other individual markers presented only 1-3 variants among the studied sample set. As these genotypes between individual loci did not always correspond, combining them provided added value for a typing approach. Two genotypes were detected for *CHS2* and *TEF1*-α, three genotypes for ITS and IGS1, and no variation was observed for Btub, except for the deviating isolates with genotype IGS1 G (Table 1, Table S1). Limited observed variation may not necessarily indicate that markers are unfit for typing but may rather reflect genetic homogeneity among deep seated isolates. Ongoing analysis of more isolates from skin and animal suggests the existence of many more genotypes than observed among the isolates in this study (B. Theelen, unpublished observation). A recently developed mating typing assay was applied to all isolates from this study, and for the *MAT b* locus only the *b1*-type was observed. Only nine isolates from one patient with IGS1-G genotype showed a somewhat deviating *MAT b* sequence. For the *MAT a* locus, both *a1* and *a2* were observed, adding value to the genetic discriminatory power of a combined marker approach. Following the previous observation of hybrid genome signatures in apparent haploid parental lineages (Theelen et al., 2022), it is relevant to mention that one of the isolates in our study contained both *MAT a1* and *MAT a2*, an observation that was also made for multiple other isolates that were not included in this study (B. Theelen, unpublished observation), further confirming the previous findings of hybrid signatures in haploid *M. furfur* strains.

### Decreased genetic variation of *M. furfur* isolates among NICU and pediatric surgery patients over the course of 5 years

Next, the determined genotype data for each individual isolate was considered per patient and compared between individuals and surveys. From the 2011 survey, 76 isolates derived from six neonatal and two pediatric patients were analyzed (Table 2, Table S1). In this group, eight unique genotypes were observed. From all but one of the neonatal patients from this group, two or more genotypes were obtained per individual. Both pediatric patients only contained one genotype, each of them unique for that specific patient (MF14 or MF16). From patient four, nine genetically deviating isolates were obtained (MF16), which was the only case of this genotype related to deep seated sources. All but three patients harbored *MAT a2b1* isolates. One deviation pertained to the unique patient four, whose isolates contained a related but deviating *MAT b1* (listed here as *MAT b1*^*^, to be further analyzed elsewhere). Pediatric patient five was the only patient with *MAT a1b1*, and from patient one, an isolate with *MAT a1a2b2* was derived, potentially pointing towards a hybrid origin, and makes this isolate an interesting candidate for induction of an *in vivo* sexual state, as Coelho and colleagues showed that a genetically constructed *MAT a1a2* transformant exhibited early stages of sexual reproduction (Coelho et al., 2023).

**Table 2.** Number of *M. furfur* isolates, genotypes, mating types and sampling sources per patient. Abbreviations: MF, *Malassezia furfur*; CVC, central venous catheter.

| Patient | # | Genotypes | Mating types | Sampling sources |
| --- | --- | --- | --- | --- |
| Survey 2011 |  |  |  |  |
| 1 | 8 | MF4, MF6, MF7, MF8 | a2b1 + a1a2b1 | Blood, CVC, skin, incubator, sheet, arm skin |
| 2 | 16 | MF4, MF8 | a2b1 | Blood, CVC, skin catheter insertion, Urine, auricular swab |
| 3 | 2 | MF8 | a2b1 | Blood, CVC |
| 4* | 9 | MF16 | a2b1* | Blood, CVC, arm skin, skin catheter insertion, blood CVC |
| 5* | 8 | MF14 | a1b1 | Blood, chest skin, skin had mother, blood CVC |
| 6 | 12 | MF4, MF5, MF8 | a2b1 | Blood, urine, CVC, conjunctival swab, gastric aspirates, conjunctival swab |
| 7 | 9 | MF4, MF8, MF12 | a2b1 | Blood, urine, chest skin, arm skin, gastric aspirates, conjunctival swab |
| 8 | 12 | MF4, MF8 | a2b1 | Blood, urine, chest skin, arm skin, gastric aspirates, conjunctival swab, endotracheal tube |
| Survey 2016 |  |  |  |  |
| 9 | 1 | MF8 | a2b1 | CVC |
| 10 | 22 | MF8 | a2b1 | Blood, Peritoneal fluid, Peritoneal catheter, TET, chest swabs, arm swabs, CVC |
| 11 | 7 | MF8 | a2b1 | Pharyngeal swab, chest swabs, arm swabs |
| 12 | 1 | MF8 | a2b1 | CVC |
| 13 | 8 | MF11, *M. slooffiae* | a2b1 | Ear swabs |
| 14 | 5 | MF8 | a2b1 | CVC, TET |
| 15 | 1 | MF8 | a2b1 | Ocular swab |
| 16 | 2 | MF12 | a1b1 | CVC |
| 17 | 1 | MF8 | a2b1 | CVC |
| 18 | 1 | MF8 | a2b1 | Ear swab |
| Reference isolates (various patients and geographies) |  |  |  |  |
| various | 13 | MF1, MF2, MF3, MF9, MF10, MF15 | a1b1 + a2b1 | Urine, blood, catheter, bronchial wash, urine, feces, nasal smear |
|  |  |  |  | France, Canada, Germany, Japan |

\* Patients 4 and 5 are 2 and 15 years old respectively and visited the Pediatric surgery ward

From the 2016 survey, 49 isolates from ten neonatal patients were obtained (table 2, Table S1). With the notion that for five patients only one isolate was obtained, this group showed a remarkably lower number of unique genotypes (MF8, MF11, MF12). Genotype MF8 was the dominant genotype with eight out of ten patients from this group only containing this variant. From patient 16, genotype MF12 was observed, a genotype that also occurred once for one patient in the 2011 survey (patient 7). From patient 13, only ear swab samples were obtained. Four out of eight isolates were identified as *Malassezia slooffiae*, and the other four isolates determined as unique genotype MF11, possibly a genotype adapted to the specific physiological conditions of the ear, or possibly a patient-specific genotype.

Comparing the observed genetic variation between both surveys, the 2011 patient group clearly was genotypically more diverse, compared to the 2016 survey (Table 2). This could possibly result from different transmission routes, multiple vs. limited points of entry into the hospital, or even survival of genotypes MF8 and MF12 over the course of five years after introduction before or during the 2011 survey. Insufficient data is available to draw conclusions about this, and extensive and continuous sampling of hospital staff and equipment is required to find clues about possible transmission routes.

In a number of cases, multiple patient sites were sampled, including skin (Table S1). For neonatal patient one the incubator and bed sheet were also sampled, and for pediatric patient five the hand of the mother was sampled. When considering genotypic variation between isolates per patient, a few observations can be made. While not a unique genotype for this sampling source, all 15 isolates derived from urine (all from the 2011 survey) retained only genotype MF4. Whether this genotype is preferential in the urine environment needs further exploration, e.g. in experiments using an animal model. Of 24 blood isolates derived from both surveys, genotypes MF4, MF8, MF14, and MF16 were observed with only genotype MF8 derived from blood of the 2016 patients, in line with the general observation that less genetic diversity is present among this patient group.

Not only was the general genetic diversity among all 2016 isolates much more limited, but every sampled patient from this group only contained only one genotype per patient, regardless of the sampling site. The only exception is patient 13 for which only ear swabs were considered and besides unique genotype MF 11 also *M. slooffiae* was found (Table 2, Table S1).

In cases where both skin and deep-seated sources were sampled (nine patients), the genotype retained from the deep-seated source was also always identified on skin, with three exceptions (Table S1). From the skin of patient one, genotypes MF4, MF6, and MF7 were observed, whereas from deep seated sources, MF4 or MF8 was obtained. For patient six, skin sampling resulted in genotypes MF4 and MF5, whereas deeper sampling sites resulted in genotypes MF4 and MF8. Finally, for patient eight, skin isolates were genotype MF4, while deep seated sampling resulted in genotypes MF4 and MF8. A few conclusions can be derived from this: 1) Almost always, the encountered genotype from deep seated sampling is also present on skin, illustrating the likely skin colonization prior to catheter-mediated transmission to deeper body site; 2) in some cases additional genotypes were observed on skin (MF6 and MF7 for patient one, and MF5 for patient six), yet deep seated environments seemed to select for genotypes MF4 and MF8). In some cases, genotype MF 8 was identified from deep seated sources while not detected from skin. This could be resulting from a sampling bias (e.g. MF8 was present on skin but missed with sampling), or a consequence of *in vivo* mutations leading to the transition from MF4 to MF8, which based on available data, would implicate mutations occuring in *IGS1* (1 SNP), *ITS* (2 SNPs + 1 indel), and *TEF*1-α (2 SNPs).

Comparing the data of both Italian surveys with the 13 reference isolates, it is interesting to note that none of the observed deep-seated genotypes from both Italian surveys were detected among the reference strains (Table 2). At the same time, none of the reference genotypes, were observed among the 125 Italian isolates, possibly suggesting the presence of unique, location or transmission source related genotypes; or specific mutations occurring at different moments, leading to unique genotypes, which is not an unlikely scenario considering that the difference between some genotypes are only one or few SNPs and/or indels.

### Exploring antifungal susceptibility trends in relationship to genotype

Where available, antifungal susceptibility test (AFST) data was assessed in relation to genotype, patient, and survey period with the notion that for only four out of eight patients from the 2011 survey AFST data was obtained (Table S1). As was previously described by Rhimi *et al*. for 39 neonatal isolates combined, low susceptibilities were observed for fluconazole (FLZ) and amphotericin B (AmB) with CLSI-based minimal inhibitory concentration (MIC) values ranging between 32-128 µg/mL after 48 hours for FLZ, and 4–32 µg/mL after 48 hours for AmB. Among the isolates considered in current study, MIC values varied between 8-128 µg/mL (mean = 90.87, sd = 42.43) for FLZ, and 0.08-16 µg/mL (mean = 7.33, sd = 5.13) for AmB. When considering MIC values for individual isolates for each of the surveys, no correlation could be found between MIC and sampling time, source, and no genotype related MIC-range was observed for FLZ or AmB. MIC values for posaconazole (POS) ranged between 0.016 and 8 µg/mL (mean = 0.41, sd = 1.23), for voriconazole (VOR) between 0.016 and 8 µg/mL (mean = 1.36, sd = 1.41), and for itraconazole (ITZ) between 0.016 and 8 µg/mL (mean = 0.39, sd = 0.06), with no observed correlation between MIC and sampling time, source, or genotype. Comparing MIC values of samples from 2011 vs 2016, for all tested antifungals, a higher mean MIC was observed for 2011, but with a high level of variation among samples (Table 3). For the two pediatric patients (2011), a higher mean MIC for VOR and ITZ was observed, compared to the neonatal patients: 3.42 (sd = 2.8) and 4.28 (sd = 7.81) vs 1.04 (sd = 0.67) and 0.16 (sd = 0.18), respectively, although for both pediatric patients, isolates with both lower and higher MIC-values were obtained. The fact that, unrelated to genotype, sampling time or source, isolates with variable MICs were observed, suggests that individual *M. furfur* isolates can adapt their susceptibility to various antifungals, independent of the observed genetic variation. Whether this is an *in vivo* exposure response or a result of an intrinsic strain-level genetic resistance mechanism, needs further exploration. Leong and colleagues observed strain-level differences in *CYP51*, a gene in which mutations have been linked with azole resistance. Their study included isolates MAL26 and MAL32 with a Y67F mutation that was not present in isolates MAL18, MAL24, or MAL25 (all MAL isolates are part of current study). This mutation seemed to insufficiently reflect differential MIC-values. Applying differential gene analysis post azole-exposure, they found an increased expression of drug transporter gene *PDR10* (Leong et al. 2021). Similar approaches for all samples from the current study might help explain the observed strain-level antifungal susceptibility variation.

**Table 3.** Mean MIC values for *Malassezia furfur* isolates for five tested antifungals for both survey years (2011 and 2016) (Iatta, Cafarchia, et al., 2014; Iatta et al., 2018; Iatta, Figueredo, et al., 2014). Abbreviations: FLZ, fluconazole; AmB, amphotericin B; POS, posaconazole; VOR, voriconazole; ITZ, itraconazole.

|  |  | FLZ | AmB | POS | VOR | ITZ |
| --- | --- | --- | --- | --- | --- | --- |
| overall | mean | 90,87 | 7,33 | 0,41 | 1,36 | 0,39 |
|  | sd | 42,44 | 5,13 | 1,23 | 1,41 | 0,06 |
| 2016 | mean | 87,00 | 4,45 | 0,06 | 0,71 | 0,09 |
|  | sd | 40,83 | 4,82 | 0,01 | 0,29 | 0,20 |
| 2011 | mean | 92,54 | 8,03 | 0,59 | 1,67 | 0,58 |
|  | sd | 43,28 | 4,99 | 1,49 | 1,62 | 2,36 |

Multiple other studies have reported reduced susceptibility for FLZ, and sometimes other azoles, and AmB (Rojas et al. 2014; Rojas et al. 2017; Gangneux et al. 2019; Leong et al. 2021; Rhimi et al. 2022) for *M. furfur* isolates, and in addition also for other *Malassezia* species (Rojas et al., 2014, 2017) and *Malassezia pachydermatis* in particular (Cafarchia et al., 2012; Álvarez-Pérez et al., 2019; Kano et al., 2019; Peano et al., 2020). Some of the reference isolates included in our study were assessed elsewhere for some of the here considered antifungals (PM315, CBS8735, CBS 14139, and CBS14141), showing variable levels of susceptibility for FLZ, AmB, and other azoles (not considered here). Although the results of some of these studies may be difficult to compare due to the lack of standardization and use of different methodologies (Rhimi, Theelen, et al., 2020) these data combined illustrate the potential for development of antifungal resistance of *M. furfur* and highlight the need for standardization and AFST monitoring.

When considering virulence, Rhimi and colleagues compared *M. furfur* isolates from PV patients and preterm infants with bloodstream infections (BSIs). They found that *M. furfur* strains from PV and BSI can produce hydrolytic enzymes. Hemolysin, lipase and phospholipase activities were lower for isolates from BSIs, whereas biofilm production was higher for those isolates (Rhimi et al., 2022). Another study compared isolates from PV and healthy subjects, and found that most strains from both groups were able to produce phospholipase, lipase, and hemolysin. When comparing mean enzymatic activities between the groups, no difference for phospholipase was found, whereas lipase activity was lower for isolates from healthy individuals, but hemolysin activity was higher (Chebil et al., 2022). The results of both studies suggest that the assessed virulence factors may play a role in pathogenesis but variation between tested groups may be dependent on external factors, rather than inherent strain differences.

Taken together, our study showed that all strains isolated from skin and deep seated neonatal or pediatric sites belong to IGS1-genotype A, except for nine isolates from one pediatric patient. This confirms previous observations of a preferential genotype for deep seated body sites. The mitochondrial *cox*3-*nad*3 intergenic marker provided additional typing resolution, and by adding ITS, *TEF1*-α, *CHS2*, and mating type, 16 overall genotypes (MF1-16) could be distinguished among all isolates from the surveys and reference isolates. These results present a promising MLST scheme that requires further testing. We observed a decreased genetic variation of *M. furfur* isolates among NICU and pediatric surgery patients over the course of 5 years and showed that the skin likely serves as a reservoir for resident *M. furfur* strains to enter the bloodstream, causing infections among susceptible patient groups. Overall, reduced susceptibility for AmB and specifically FLZ was observed.

## Materials and Methods

### Strains, media, and growth conditions

A total of 125 isolates were obtained from two surveys done in Italian NICU and pediatric wards, and stored in glycerol stocks at -80 °C until further processing. All isolates from the Italian 2011 (n = 76) and 2016 (n = 49) surveys, and 13 additional reference strains were revived from glycerol stocks on MLNA petri dishes for 72 h at 30 °C, after which a subculture was incubated on mDixon medium for 48 h at 30 °C, prior to DNA-extraction.

### DNA extraction

DNA was extracted according to the CTAB method as described by O’Donnell et al. (O’Donnell et al., 1997), with some modifications: two 10 μl loops of fresh biomass were suspended in 750 μl CTAB buffer and mechanically disrupted in a TissueLyser II (Qiagen®) at 30Hz for 8 min.The mixture was heated for 1 h at 65 °C with frequent resuspension using a vortex. After lysis, two purification steps were performed with phenol– chloroform and chloroform, respectively. Extracted DNA was dissolved in 100 μl TE and 10-50x diluted in MQ water.

### PCR reactions, sequencing and phylogenetic analysis

PCR and sequencing reactions for the *MAT a* and *MAT b* loci, as well as the ITS, IGS1, and protein coding genes *TEF*1-α and *CHS2* were performed as previously described (Bart Theelen et al. 2022). Raw sequence data were evaluated and consensus sequences created with Seqman ProTM (v. 9.0.4 (39), DNASTAR, Inc., Madison, USA). Sequence alignments and phylogenetic analyses were conducted in Geneious Prime, version 2022.0.1 (Biomatters Ltd.). Sequences for each individual locus were aligned with Muscle, version 3.8.425 with following settings: kmer4_6 and pctid_kimura for distance measure, UPGMB for all clustering method interactions, “pseudo” for tree rooting method, and ClustalW for the sequence weighting scheme (Edgar 2004). Phylogenetic trees were constructed with the FastTree version 2.1.11 plugin, using standard settings (Price, Dehal, and Arkin 2010) with additional formatting using FigTree, version 1.4.4(https://github.com/rambaut/figtree/releases/tag/v1.4.4).

### Antifungal susceptibility testing

Part of the MIC-data has partially been published previously in a grouped manner. MIC-values were determined as previously described (Iatta et al., 2014; Rhimi et al., 2020). Here, data obtained with the Clinical and Laboratory Standards Institute (CLSI) method after 48 h of incubation was used.

## Acknowledgements

We thank Koichi Makimura for providing the isolates GIFU01 and GIFU02

