## Supplementary table S1 for "Limited genetic diversity among *Malassezia furfur* isolates during two one-year neonatal and pediatric bloodstream infection surveys confirms preference for one specific IGS1 genotype in deep-seated infections"

**Table S1**. Overview of survey isolates and reference strains, with sample backgrounds and genotyping, mating typing data and MIC values when tested. Abbreviations: MF, *Malassezia furfur*; IGS1, intergenic spacer 1 region; ITS, internal transcribed spacer; Btub, ϐ-tubulin; CHS2, chitin synthase 2; *TEF1*-α, translation elongation factor 1-α; *cox*3-*nad*3, *cox*3-*nad*3 intergenic region; CLSI, Clinical and Laboratory Standards Institute; CVC, central venous catheter; TET, Endotracheal tube; NICU, neonatal intensive care unit.

|  |  |  |  |  |  |  |  | | | | |  | **Mating type** | | | | | **AFST (CLSI)** | | | | |
| --- | --- | --- | --- | --- | --- | --- | --- | --- | --- | --- | --- | --- | --- | --- | --- | --- | --- | --- | --- | --- | --- | --- |
| **strain** | **source** | **location** | **#** | **age** | **gender** | **MF** | **IGS** | **ITS** | ***TEF1*-α** | ***CH2S*** | **Btub** | ***cox*3-*nad*3** | **a1** | **a2** | **b1b3** | | **b2b4** | **POS** | **VORI** | **ITZ** | **FLZ** | **AMB** |
| CBS14139 | Urine, human | France | NA | NA | NA | MF10 | A2 | N | V | II | µ | l8 | 1 | 0 | | 1 | 0 |  |  |  |  |  |
| CBS14140 | Catheter, blood | France | NA | NA | NA | MF9 | A2 | N | V | II | µ | l4 | 0 | 1 | | 1 | 0 |  |  |  |  |  |
| CBS14141 | Human,catheter, blood | France | NA | NA | NA | MF9 | A2 | N | V | II | µ | l4 | 0 | 1 | | 1 | 0 |  |  |  |  |  |
| CBS8735 | Bronchial wash, man | Canada | NA | NA | NA | MF2 | A1 | K | V | II | µ | l2 | 1 | 0 | | 1 | 0 |  |  |  |  |  |
| PM312 | Urine of neonate | Germany | NA | NA | NA | MF3 | A1 | K | V | II | µ | l3 | 1 | 0 | | 1 | 0 |  |  |  |  |  |
| PM315 | Anal swab of neonate | Germany | NA | NA | NA | MF3 | A1 | K | V | II | µ | l3 | 1 | 0 | | 1 | 0 |  |  |  |  |  |
| PM317 | Feces of neonate | Germany | NA | NA | NA | MF3 | A1 | K | V | II | µ | l3 | 1 | 0 | | 1 | 0 |  |  |  |  |  |
| PM318 | Feces of neonate | Germany | NA | NA | NA | MF3 | A1 | K | V | II | µ | l3 | 1 | 0 | | 1 | 0 |  |  |  |  |  |
| PM319 | Nasal smear of neonate | Germany | NA | NA | NA | MF3 | A1 | K | V | II | µ | l3 | 1 | 0 | | 1 | 0 |  |  |  |  |  |
| GIFU01 | Blood | Japan | NA | NA | NA | MF15 | A2 | N | W | III | µ | i3 | 1 | 0 | | 1 | 0 |  |  |  |  |  |
| GIFU02 | Catheter tip | Japan | NA | NA | NA | MF15 | A2 | N | W | III | µ | i3 | 1 | 0 | | 1 | 0 |  |  |  |  |  |
| CBS7983 | Blood of leukemia patient | France | NA | NA | NA | MF1 | A1 | K | V | II | µ | i4 | 1 | 0 | | 1 | 0 |  |  |  |  |  |
| CBS9767 | Catheter, blood, liver recipient | France | NA | NA | NA | MF9 | A2 | N | V | II | µ | l4 | 0 | 1 | | 1 | 0 |  |  |  |  |  |
| CD1465 | CVC | NICU | 9 |  |  | MF8 | A2 | N | V | II | µ | i4 | 0 | 1 | | 1 | 0 |  |  |  |  |  |
| CD1472 | Blood | NICU | 10 |  |  | MF8 | A2 | N | V | II | µ | i4 | 0 | 1 | | 1 | 0 | 0,06 | 1 | 0,125 | >128 | 8 |
| CD1473 | Peritoneal fluid | NICU | 10 |  |  | MF8 | A2 | N | V | II | µ | i4 | 0 | 1 | | 1 | 0 | 0,06 | 1 | 0,125 | >128 | 8 |
| CD1474 | Peritoneal catheter | NICU | 10 |  |  | MF8 | A2 | N | V | II | µ | i4 | 0 | 1 | | 1 | 0 | 0,06 | 1 | 0,008 | >128 | 0,008 |
| CD1475 | TET | NICU | 10 |  |  | MF8 | A2 | N | V | II | µ | i4 | 0 | 1 | | 1 | 0 | 0,016 | 1 | 0,06 | >128 | 0,125 |
| CD1476 | Chest swab (colony 1) | NICU | 10 |  |  | MF8 | A2 | N | V | II | µ | i4 | 0 | 1 | | 1 | 0 | 0,06 | 1 | 0,06 | >128 | 4 |
| CD1477 | Chest swab (colony 2) | NICU | 10 |  |  | MF8 | A2 | N | V | II | µ | i4 | 0 | 1 | | 1 | 0 | 0,06 | 1 | 0,008 | >128 | 0,008 |
| CD1478 | Chest swab (colony 3) | NICU | 10 |  |  | MF8 | A2 | N | V | II | µ | i4 | 0 | 1 | | 1 | 0 | 0,06 | 1 | 0,008 | >128 | 0,008 |
| CD1479 | Chest swab (colony 4) | NICU | 10 |  |  | MF8 | A2 | N | V | II | µ | i4 | 0 | 1 | | 1 | 0 | 0,06 | 0,5 | 0.016 | >128 | 2 |
| CD1480 | Chest swab (colony 5) | NICU | 10 |  |  | MF8 | A2 | N | V | II | µ | i4 | 0 | 1 | | 1 | 0 | 0,06 | 0,25 | 0.016 | >128 | 2 |
| CD1481 | Chest swab (colony 6) | NICU | 10 |  |  | MF8 | A2 | N | V | II | µ | i4 | 0 | 1 | | 1 | 0 | 0,06 | 0,016 | 0.06 | >128 | 4 |
| CD1482 | Chest swab (colony 7) | NICU | 10 |  |  | MF8 | A2 | N | V | II | µ | i4 | 0 | 1 | | 1 | 0 | 0,06 | 0,5 | 0,008 | 32 | >16 |
| CD1483 | Chest swab (colony 8) | NICU | 10 |  |  | MF8 | A2 | N | V | II | µ | i4 | 0 | 1 | | 1 | 0 | 0,125 | 1 | 0,06 | 64 | 4 |
| CD1484 | Chest swab (colony 9) | NICU | 10 |  |  | MF8 | A2 | N | V | II | µ | i4 | 0 | 1 | | 1 | 0 | 0,06 | 0,5 | 0,03 | 32 | 2 |
| CD1485 | Arm swab (colony 1) | NICU | 10 |  |  | MF8 | A2 | N | V | II | µ | i4 | 0 | 1 | | 1 | 0 | 0,06 | 1 | 0,06 | >128 | 4 |
| CD1486 | Arm swab (colony 2) | NICU | 10 |  |  | MF8 | A2 | N | V | II | µ | i4 | 0 | 1 | | 1 | 0 |  |  |  |  |  |
| CD1487 | Arm swab (colony 3) | NICU | 10 |  |  | MF8 | A2 | N | V | II | µ | i4 | 0 | 1 | | 1 | 0 |  |  |  |  |  |
| CD1488 | Arm swab (colony 4) | NICU | 10 |  |  | MF8 | A2 | N | V | II | µ | i4 | 0 | 1 | | 1 | 0 | 0,06 | 1 | 0,06 | 64 | >16 |
| CD1489 | Arm swab (colony 5) | NICU | 10 |  |  | MF8 | A2 | N | V | II | µ | i4 | 0 | 1 | | 1 | 0 | 0,06 | 1 | 0,06 | 64 |  |
| CD1490 | Arm swab (colony 6) | NICU | 10 |  |  | MF8 | A2 | N | V | II | µ | i4 | 0 | 1 | | 1 | 0 | 0,06 | 1 | 0,06 | 64 |  |
| CD1491 | Arm swab (colony 7) | NICU | 10 |  |  | MF8 | A2 | N | V | II | µ | i4 | 0 | 1 | | 1 | 0 | 0,06 | 1 | 0,06 | 64 |  |
| CD1492 | Arm swab (colony 8) | NICU | 10 |  |  | MF8 | A2 | N | V | II | µ | i4 | 0 | 1 | | 1 | 0 | 0,06 | 1 | 0,06 | 64 |  |
| CD1493 | CVC | NICU | 10 |  |  | MF8 | A2 | N | V | II | µ | i4 | 0 | 1 | | 1 | 0 | 0,06 | 0,5 | 1 | 128 | 4 |
| CD1494 | Pharyngeal swab | NICU | 11 |  |  | MF8 | A2 | N | V | II | µ | i4 | 0 | 1 | | 1 | 0 | 0,06 | 0,5 | 0,03 | 32 |  |
| CD1515 | Chest swab (colony 1) | NICU | 11 |  |  | MF8 | A2 | N | V | II | µ | i4 | 0 | 1 | | 1 | 0 | 0,06 | 1 | 0,03 | >128 | 2 |
| CD1516 | Chest swab (colony 2) | NICU | 11 |  |  | MF8 | A2 | N | V | II | µ | i4 | 0 | 1 | | 1 | 0 | 0,06 | 0,5 | 0,03 | 32 |  |
| CD1517 | Chest swab (colony 3) | NICU | 11 |  |  | MF8 | A2 | N | V | II | µ | i4 | 0 | 1 | | 1 | 0 | 0,06 | 0,5 | 0,016 | 32 |  |
| CD1518 | Arm swab (colony 1) | NICU | 11 |  |  | MF8 | A2 | N | V | II | µ | i4 | 0 | 1 | | 1 | 0 | 0,06 | 0,5 | 0,03 | 64 |  |
| CD1519 | Arm swab (colony 2) | NICU | 11 |  |  | MF8 | A2 | N | V | II | µ | i4 | 0 | 1 | | 1 | 0 | 0,06 | 0,5 | 0,03 | 64 |  |
| CD1520 | Arm swab (col.3) | NICU | 11 |  |  | MF8 | A2 | N | V | II | µ | i4 | 0 | 1 | | 1 | 0 | 0,06 | 0,5 | 0,03 | 64 |  |
| CD1495 | CVC | NICU | 12 |  | F | MF8 | A2 | N | V | II | µ | i4 | 0 | 1 | | 1 | 0 | 0,06 | 0,5 | 0,5 | 128 | 4 |
| CD1510 | Ear swab (colony 2 rugosa) | NICU | 13 |  |  | MF11 | A2 | N | W | II | µ | i2 | 0 | 1 | | 1 | 0 | 0,06 | 0,5 | 0,03 | 32 |  |
| CD1511 | Ear swab (colony 3 rugosa) | NICU | 13 |  |  | *M*. slooffiae | NA | NA | NA | NA | NA | NA | NA | NA | | NA | NA | NA | NA | NA | NA |  |
| CD1512 | Ear swab (colony 1 piccola) | NICU | 13 |  |  | *M*. slooffiae | NA | NA | NA | NA | NA | NA | NA | NA | | NA | NA | NA | NA | NA | NA |  |
| CD1513 | Ear swab (colony 2 piccola) | NICU | 13 |  |  | *M*. slooffiae | NA | NA | NA | NA | NA | NA | NA | NA | | NA | NA | NA | NA | NA | NA |  |
| CD1514 | Ear swab (colony 3 piccola) | NICU | 13 |  |  | *M*. slooffiae | NA | NA | NA | NA | NA | NA | NA | NA | | NA | NA | SLOOF | SLOOF | SLOOF | SLOOF |  |
| CD1521 | Ear swab (colony 1) | NICU | 13 |  |  | MF11 | A2 | N | W | II | µ | i2 | 0 | 1 | | 1 | 0 | 0,06 | 0,5 | 0,03 | 128 |  |
| CD1522 | Ear swab (colony 2) | NICU | 13 |  |  | MF11 | A2 | N | W | II | µ | i2 | 0 | 1 | | 1 | 0 | 0,06 | 0,5 | 0,03 | 64 |  |
| CD1523 | Ear swab (colony 3) | NICU | 13 |  |  | MF11 | A2 | N | W | II | µ | i2 | 0 | 1 | | 1 | 0 | 0,06 | 0,5 | 0,03 | 32 |  |
| CD1530 | CVC (colony 1) | NICU | 14 |  |  | MF8 | A2 | N | V | II | µ | i4 | 0 | 1 | | 1 | 0 |  |  |  |  |  |
| CD1531 | TET (colony 2) | NICU | 14 |  |  | MF8 | A2 | N | V | II | µ | i4 | 0 | 1 | | 1 | 0 |  |  |  |  |  |
| CD1532 | TET (colony 1) | NICU | 14 |  |  | MF8 | A2 | N | V | II | µ | i4 | 0 | 1 | | 1 | 0 |  |  |  |  |  |
| CD1533 | TET (colony 2) | NICU | 14 |  |  | MF8 | A2 | N | V | II | µ | i4 | 0 | 1 | | 1 | 0 |  |  |  |  |  |
| CD1534 | TET (colony 3) | NICU | 14 |  |  | MF8 | A2 | N | V | II | µ | i4 | 0 | 1 | | 1 | 0 |  |  |  |  |  |
| CD1545 | Ocular swab | NICU | 15 |  |  | MF8 | A2 | N | V | II | µ | i4 | 0 | 1 | | 1 | 0 |  |  |  |  |  |
| CD1547 | CVC | NICU | 16 |  |  | MF12 | A2 | N | W | II | µ | i4 | 1 | 0 | | 1 | 0 |  |  |  |  |  |
| CD1548 | CVC | NICU | 16 |  |  | MF12 | A2 | N | W | II | µ | i4 | 1 | 0 | | 1 | 0 |  |  |  |  |  |
| CD1549 | CVC | NICU | 17 |  | M | MF8 | A2 | N | V | II | µ | i4 | 0 | 1 | | 1 | 0 |  |  |  |  |  |
| CD1559 | Ear swab | ICU | 18 |  | M | MF8 | A2 | N | V | II | µ | i4 | 0 | 1 | | 1 | 0 |  |  |  |  |  |
| MAL01 | Blood | NICU | 1 | 2 m | M | MF4 | A1 | K | W | II | µ | i4 | 0 | 1 | | 1 | 0 | 0,06 | 0,06 | 0,03 | 16 | 8 |
| MAL02 | Blood | NICU | 1 | 3 m | M | MF8 | A2 | N | V | II | µ | i4 | 0 | 1 | | 1 | 0 | 0,03 | 0,25 | 0,03 | 128 | 4 |
| MAL03 | CVC | NICU | 1 | 3 m | M | MF4 | A1 | K | W | II | µ | i4 | 0 | 1 | | 1 | 0 | 0,25 | 0,5 | 0,25 | 128 | 8 |
| MAL04 | Skin | NICU | 1 | 3 m | M | MF4 | A1 | K | W | II | µ | i4 | 0 | 1 | | 1 | 0 | 0,125 | 2 | 0,125 | 128 | 8 |
| MAL05 | Incubator | NICU | 1 | 3 m | M | MF4 | A1 | K | W | II | µ | i4 | 0 | 1 | | 1 | 0 |  |  |  |  |  |
| MAL06 | Sheet | NICU | 1 | 3 m | M | MF4 | A1 | K | W | II | µ | i4 | 0 | 1 | | 1 | 0 |  |  |  |  |  |
| MAL07 | Arm skin | NICU | 1 | 3 m | M | MF6 | A1 | N | W | II | µ | i4 | 0 | 1 | | 1 | 0 | 0,06 | 2 | 0,06 | 128 | 8 |
| MAL09 | Arm skin | NICU | 1 | 3 m | M | MF7 | A2 | N | V | II | µ | i4 | 1 | 1 | | 1 | 0 | 0,25 | 2 | 0,25 | 128 | 8 |
| MAL11 | Blood | NICU | 2 | 3 m | M | MF8 | A2 | N | V | II | µ | i4 | 0 | 1 | | 1 | 0 | 0,125 | 1 | 0,125 | 16 | 16 |
| MAL12 | Blood | NICU | 2 | 3 m | M | MF8 | A2 | N | V | II | µ | i4 | 0 | 1 | | 1 | 0 | 1 | 1 | 1 | 32 | 16 |
| MAL13 | Blood | NICU | 2 | 3 m | M | MF8 | A2 | N | V | II | µ | i4 | 0 | 1 | | 1 | 0 | 0,125 | 0,5 | 0,25 | 32 | 16 |
| MAL14 | CVC | NICU | 2 | 3 m | M | MF8 | A2 | N | V | II | µ | i4 | 0 | 1 | | 1 | 0 | 0,5 | 0,5 | 0,25 | 32 | 16 |
| MAL15 | Skin catheter insertion | NICU | 2 | 3 m | M | MF8 | A2 | N | V | II | µ | i4 | 0 | 1 | | 1 | 0 | 0,06 | 0,5 | 0,06 | 32 | 8 |
| MAL16 | Skin catheter insertion | NICU | 2 | 3 m | M | MF8 | A2 | N | V | II | µ | i4 | 0 | 1 | | 1 | 0 | 0,06 | 1 | 0,06 | 16 | 4 |
| MAL26 | Blood | NICU | 2 | 4 m | M | MF4 | A1 | K | W | II | µ | i4 | 0 | 1 | | 1 | 0 | 0,25 | 1 | 0,25 | 128 | 8 |
| MAL27 | Blood | NICU | 2 | 4 m | M | MF4 | A1 | K | W | II | µ | i4 | 0 | 1 | | 1 | 0 | 0,25 | 2 | 0,25 | 128 | 8 |
| MAL28 | Skin catheter insertion | NICU | 2 | 4 m | M | MF4 | A1 | K | W | II | µ | i4 | 0 | 1 | | 1 | 0 | 0,25 | 1 | 0,25 | 128 | 8 |
| MAL29 | Urine | NICU | 2 | 4 m | M | MF4 | A1 | K | W | II | µ | i4 | 0 | 1 | | 1 | 0 | 0,25 | 1 | 0,25 | 128 | 4 |
| MAL30 | Urine | NICU | 2 | 4 m | M | MF4 | A1 | K | W | II | µ | i4 | 0 | 1 | | 1 | 0 | 0,25 | 1 | 0,25 | 128 | 4 |
| MAL31 | Urine | NICU | 2 | 5 m | M | MF4 | A1 | K | W | II | µ | i4 | 0 | 1 | | 1 | 0 | 0,25 | 1 | 0,25 | 128 | 8 |
| MAL32 | Urine | NICU | 2 | 5 m | M | MF4 | A1 | K | W | II | µ | i4 | 0 | 1 | | 1 | 0 | 2 | 4 | 4 | 128 | 8 |
| MAL33 | Urine | NICU | 2 | 5 m | M | MF4 | A1 | K | W | II | µ | i4 | 0 | 1 | | 1 | 0 | 0,25 | 2 | 0,25 | 128 | 16 |
| MAL34 | Urine | NICU | 2 | 5 m | M | MF4 | A1 | K | W | II | µ | i4 | 0 | 1 | | 1 | 0 | 0,25 | 1 | 0,25 | 128 | 16 |
| MAL35 | Auricolar swab | NICU | 2 | 6 m | M | MF8 | A2 | N | V | II | µ | i4 | 0 | 1 | | 1 | 0 | 0,125 | 0,5 | 0,125 | 64 | 8 |
| MAL39 | Blood from CVC | NICU | 3 | 1 m | F | MF8 | A2 | N | V | II | µ | i4 | 0 | 1 | | 1 | 0 | 0,125 | 0,5 | 0,5 | 32 | 8 |
| MAL40 | CVC | NICU | 3 | 1 m | F | MF8 | A2 | N | V | II | µ | i4 | 0 | 1 | | 1 | 0 | 1 | 1 | 0,125 | 64 | 16 |
| MAL17 | Blood from CVC | Pediatric surgery | 4 | 2 y | M | MF16 | G1 | L | W | III | ∆2 | l7 | 0 | 1 | | **1** | 0 | 0,016 | 0,06 | 0,03 | 128 | 1 |
| MAL18 | Blood | Pediatric surgery | 4 | 2 y | M | MF16 | G1 | L | W | III | ∆2 | l7 | 0 | 1 | | **1** | 0 | 2 | 0,125 | 8 | 128 | 8 |
| MAL19 | Blood | Pediatric surgery | 4 | 2 y | M | MF16 | G1 | L | W | III | ∆2 | l7 | 0 | 1 | | **1** | 0 | 0,125 | 0,5 | 0,125 | 16 | 16 |
| MAL20 | Blood | Pediatric surgery | 4 | 2 y | M | MF16 | G1 | L | W | III | ∆2 | l7 | 0 | 1 | | **1** | 0 | 8 | 8 | 8 | 64 | 16 |
| MAL21 | Blood from CVC (Broviack) | Pediatric surgery | 4 | 2 y | M | MF16 | G1 | L | W | III | ∆2 | l7 | 0 | 1 | | **1** | 0 | 0,125 | 1 | 0,25 | 32 | 8 |
| MAL22 | Blood from CVC (Broviack) | Pediatric surgery | 4 | 2 y | M | MF16 | G1 | L | W | III | ∆2 | l7 | 0 | 1 | | **1** | 0 | 0,25 | 2 | 0,5 | 128 | 8 |
| MAL23 | Blood from CVC (Broviack) | Pediatric surgery | 4 | 2 y | M | MF16 | G1 | L | W | III | ∆2 | l7 | 0 | 1 | | **1** | 0 | 1 | 2 | 4 | 32 | 8 |
| MAL24 | Arm skin | Pediatric surgery | 4 | 2 y | M | MF16 | G1 | L | W | III | ∆2 | l7 | 0 | 1 | | **1** | 0 | 8 | 8 | 16 | 128 | 16 |
| MAL25 | Skin catheter insertion | Pediatric surgery | 4 | 2 y | M | MF16 | G1 | L | W | III | ∆2 | l7 | 0 | 1 | | **1** | 0 | 0,25 | 1 | 0,25 | 128 | 16 |
| MAL36 | Blood | Pediatric surgery | 5 | 15 y | F | MF14 | A2 | N | W | II | µ | l1 | 1 | 0 | | 1 | 0 | 0,25 | 0,5 | 0,125 | 64 | 8 |
| MAL37 | Chest skin | Pediatric surgery | 5 | 15 y | F | MF14 | A2 | N | W | II | µ | l1 | 1 | 0 | | 1 | 0 | 0,25 | 8 | 0,5 | 128 | 2 |
| MAL38 | Mother hand skin | Pediatric surgery | 5 | 15 y | F | MF14 | A2 | N | W | II | µ | l1 | 1 | 0 | | 1 | 0 | 0,25 | 2 | 0,5 | 128 | 8 |
| MAL41 | Blood from CVC | Pediatric surgery | 5 | 15 y | F | MF14 | A2 | N | W | II | µ | l1 | 1 | 0 | | 1 | 0 | 0,125 | 2 | 0,25 | 128 | 1 |
| MAL42 | Blood | Pediatric surgery | 5 | 15 y | F | MF14 | A2 | N | W | II | µ | l1 | 1 | 0 | | 1 | 0 | 0,25 | 4 | 0,5 | 128 | 16 |
| MAL43 | Blood from CVC | Pediatric surgery | 5 | 15 y | F | MF14 | A2 | N | W | II | µ | l1 | 1 | 0 | | 1 | 0 | 4 | 2 | 1 | 8 | 16 |
| MAL44 | Blood | Pediatric surgery | 5 | 15 y | F | MF14 | A2 | N | W | II | µ | l1 | 1 | 0 | | 1 | 0 | 0,25 | 4 | 0,5 | 128 | 8 |
| MAL45 | Blood from CVC | Pediatric surgery | 5 | 15 y | F | MF14 | A2 | N | W | II | µ | l1 | 1 | 0 | | 1 | 0 | 0,25 | 4 | 0,5 | 64 | 16 |
| MAL46 | Blood | NICU | 6 | 1m | M | MF8 | A2 | N | V | II | µ | i4 | 0 | 1 | | 1 | 0 | 0,06 | 0,25 | 0,03 | 64 | 4 |
| MAL48 | Conjunctival swab | NICU | 6 | 1m | M | MF5 | A1 | N | V | II | µ | i4 | 0 | 1 | | 1 | 0 | 0,125 | 0,5 | 0,125 | 64 | 8 |
| MAL49 | Gastric aspirates | NICU | 6 | 1m | M | MF4 | A1 | K | W | II | µ | i4 | 0 | 1 | | 1 | 0 | 0,25 | 2 | 0,25 | 64 | 8 |
| MAL50 | Chest skin | NICU | 6 | 1m | M | MF5 | A1 | N | V | II | µ | i4 | 0 | 1 | | 1 | 0 | 0,06 | 0,25 | 0,016 | 64 | 8 |
| MAL51 | Urine | NICU | 6 | 1m | M | MF4 | A1 | K | W | II | µ | i4 | 0 | 1 | | 1 | 0 | 0,25 | 1 | 0,25 | 64 | 8 |
| MAL52 | CVC | NICU | 6 | 1m | M | MF5 | A2 | N | V | II | µ | i4 | 0 | 1 | | 1 | 0 | 0,06 | 0,25 | 0,06 | 128 | 16 |
| MAL61 | Urine | NICU | 6 | 1m | M | MF4 | A1 | K | W | II | µ | i4 | 0 | 1 | | 1 | 0 | 0,25 | 2 | 0,25 | 64 | 1 |
| MAL62 | Conjunctival swab | NICU | 6 | 1m | M | MF4 | A1 | K | W | II | µ | i4 | 0 | 1 | | 1 | 0 | 0,125 | 1 | 0,25 | 64 | 2 |
| MAL63 | Arm skin | NICU | 6 | 1m | M | MF4 | A1 | K | W | II | µ | i4 | 0 | 1 | | 1 | 0 | 0,25 | 2 | 0,25 | 128 | 2 |
| MAL69 | Urine | NICU | 6 | 2m | M | MF4 | A1 | K | W | II | µ | i4 | 0 | 1 | | 1 | 0 | 0,25 | 2 | 0,25 | 128 | 4 |
| MAL75 | Gastric aspirates | NICU | 6 | 2m | M | MF8 | A2 | N | V | II | µ | i4 | 0 | 1 | | 1 | 0 | 0,25 | 1 | 0,25 | 128 | 8 |
| MAL77 | Urine | NICU | 6 | 2m | M | MF4 | A1 | K | W | II | µ | i4 | 0 | 1 | | 1 | 0 | 0,125 | 2 | 0,125 | 128 | 8 |
| MAL47 | Blood | NICU | 7 | 1 m | M | MF8 | A2 | N | V | II | µ | i4 | 0 | 1 | | 1 | 0 | 0,06 | 0,25 | 0,06 | 64 | 4 |
| MAL55 | Chest skin | NICU | 7 | 1 m | M | MF8 | A2 | N | V | II | µ | i4 | 0 | 1 | | 1 | 0 | 0,25 | 1 | 0,25 | 128 | 16 |
| MAL56 | Urine | NICU | 7 | 1 m | M | MF4 | A1 | K | W | II | µ | i4 | 0 | 1 | | 1 | 0 | 0,25 | 1 | 0,25 | 64 | 2 |
| MAL57 | Gastric aspirates | NICU | 7 | 1 m | M | MF12 | A2 | N | W | II | µ | i4 | 0 | 1 | | 1 | 0 | 0,25 | 1 | 0,25 | 64 | 4 |
| MAL58 | CVC | NICU | 7 | 1 m | M | MF8 | A2 | N | V | II | µ | i4 | 0 | 1 | | 1 | 0 | 0,06 | 0,25 | 0,125 | 16 | 2 |
| MAL59 | Urine | NICU | 7 | 1 m | M | MF4 | A1 | K | W | II | µ | i4 | 0 | 1 | | 1 | 0 | 0,25 | 2 | 0,25 | 64 | 2 |
| MAL64 | Arm skin | NICU | 7 | 1 m | M | MF4 | A1 | K | W | II | µ | i4 | 0 | 1 | | 1 | 0 | 0,25 | 2 | 0,25 | 128 | 16 |
| MAL65 | Conjunctival swab | NICU | 7 | 1 m | M | MF4 | A1 | K | W | II | µ | i4 | 0 | 1 | | 1 | 0 | 0,125 | 1 | 0,25 | 64 | 2 |
| MAL70 | Urine | NICU | 7 | 1 m | M | MF4 | A1 | K | W | II | µ | i4 | 0 | 1 | | 1 | 0 | 0,25 | 1 | 0,25 | 128 | 8 |
| MAL53 | Blood | NICU | 8 | 1 m | M | MF8 | A2 | N | V | II | µ | i4 | 0 | 1 | | 1 | 0 | 0,06 | 0,5 | 0,125 | 128 | 4 |
| MAL54 | Chest skin | NICU | 8 | 1 m | M | MF4 | A1 | K | W | II | µ | i4 | 0 | 1 | | 1 | 0 | 0,25 | 2 | 0,25 | 128 | 1 |
| MAL60 | Urine | NICU | 8 | 1 m | M | MF4 | A1 | K | W | II | µ | i4 | 0 | 1 | | 1 | 0 | 0,25 | 2 | 0,25 | 64 | 4 |
| MAL66 | arm skin | NICU | 8 | 1 m | M | MF4 | A1 | K | W | II | µ | i4 | 0 | 1 | | 1 | 0 | 0,25 | 2 | 0,25 | 8 | 2 |
| MAL67 | Conjunctival swab | NICU | 8 | 1 m | M | MF4 | A1 | K | W | II | µ | i4 | 0 | 1 | | 1 | 0 | 0,125 | 2 | 0,125 | 128 | 2 |
| MAL68 | gastric aspirates | NICU | 8 | 1 m | M | MF4 | A1 | K | W | II | µ | i4 | 0 | 1 | | 1 | 0 | 0,125 | 1 | 0,25 | 128 | 4 |
| MAL71 | Urine | NICU | 8 | 1 m | M | MF4 | A1 | K | W | II | µ | i4 | 0 | 1 | | 1 | 0 | 0,125 | 1 | 0,25 | 128 | 8 |
| MAL72 | Endotracheal tube | NICU | 8 | 1 m | M | MF4 | A1 | K | W | II | µ | i4 | 0 | 1 | | 1 | 0 | 0,25 | 1 | 0,25 | 128 | 4 |
| MAL73 | Urine | NICU | 8 | 1 m | M | MF4 | A1 | K | W | II | µ | i4 | 0 | 1 | | 1 | 0 | 0,25 | 2 | 0,25 | 128 | 4 |
| MAL74 | Gastric aspirates | NICU | 8 | 1 m | M | MF4 | A1 | K | W | II | µ | i4 | 0 | 1 | | 1 | 0 | 0,25 | 1 | 0,25 | 128 | 8 |
| MAL76 | Endotracheal tube | NICU | 8 | 1 m | M | MF4 | A1 | N | W | II | µ | i4 | 0 | 1 | | 1 | 0 | 0,25 | 2 | 0,25 | 128 | 8 |
| MAL78 | Gastric aspirates | NICU | 8 | 1 m | M | MF8 | A2 | N | V | II | µ | i4 | 0 | 1 | | 1 | 0 | 0,125 | 1 | 0,25 | 128 | 8 |
